# Graded Variation In Cortical T1w/T2w Myelination During Adolescence

**DOI:** 10.1101/2021.12.06.471432

**Authors:** Graham L. Baum, John C. Flournoy, Matthew F. Glasser, Michael P. Harms, Patrick Mair, Ashley Sanders, Deanna M. Barch, Randy L. Buckner, Susan Bookheimer, Mirella Dapretto, Stephen Smith, Kathleen M. Thomas, Essa Yacoub, David C. Van Essen, Leah H. Somerville

## Abstract

Myelination influences brain connectivity during sensitive periods of development by enhancing neural signaling speed and regulating synapse formation to reduce plasticity. However, *in vivo* studies characterizing the maturational timing of cortical myelination during human development remain scant. Here, we take advantage of recent advances in high-resolution cortical T1w/T2w myelin mapping methods, including principled correction of B1+ transmit field effects, using data from the Human Connectome Project in Development (*N*=628, ages 8-21) to characterize the maturational timing of myelination from childhood through early adulthood throughout the cerebral neocortex. We apply Bayesian spline models and functional latent clustering analysis to demonstrate graded variation in the rate of cortical T1w/T2w myelin growth in neocortical areas that is strongly correlated with the sensorimotor-association (S-A) axis of cortical organization reported by others. In sensorimotor areas T1w/T2w myelin starts at high levels at early ages, increases at a fast pace, and decelerates at later ages (18-21). In intermediate multimodal areas along the S-A axis, T1w/T2w myelin tends to start at intermediate levels and increase linearly at an intermediate pace. In transmodal/paralimbic association areas high along the S-A axis, T1w/T2w myelin tends to start at low levels and increase linearly at the slowest pace. These data provide evidence for graded variation along the S-A axis in the rate of cortical myelination during adolescence, which could reflect ongoing plasticity underlying the development of complex information processing and psychological functioning.

**Significance Statement:** Myelin is a lipid membrane that is essential to healthy brain function. Myelin wraps axons to increase neural signaling speed, enabling complex neuronal functioning underlying learning and cognition. Here we characterize the developmental timing of myelination across the cerebral cortex during adolescence using recent advances in non-invasive myelin mapping. Our results provide new evidence demonstrating graded variation across the cortex in the timing of myelination during adolescence, with rapid myelination in lower-order sensory areas and gradual myelination in higher-order association areas. This spatial pattern of microstructural brain development closely parallels the sensorimotor-to-association axis of cortical organization and plasticity during ontogeny.

## INTRODUCTION

Adolescence, the transition from childhood to adulthood, is characterized by the refinement and stabilization of neural circuits supporting the dynamic control of attention and behavior (Larsen and Luna, 2018). Axonal myelination in the cerebral cortex enhances signaling speed (McDougall et al., 2018) and regulates synapse formation during development to support the reliable instantiation of adaptive behavior (Makinodan et al., 2012; Mount and Monje, 2017). While the role of cortical myelination in regulating plasticity during sensitive periods of neurodevelopment has been demonstrated in animal models (Takesian and Hensch, 2013; Toyoizumi et al., 2013), there are few *in vivo* studies characterizing the maturational timing of cortical myelination during sensitive periods of human development.

Cascading sensitive periods for increasingly complex cognitive functions are associated with progressive refinement of neural systems along the sensorimotor-association (S-A) axis of cortical organization (Sydnor et al., 2021). The S-A axis represents a continuum of anatomical, functional, metabolic, and genetic variation that underpins variability in cortical information processing. Sensitive periods in the first years of postnatal development involve synaptic remodeling in sensorimotor cortex for reliable perception and movement (Takesian and Hensch, 2013). Throughout adolescence and early adulthood, sensitive periods for developing cognitive control and socioemotional processing may reflect the protracted refinement of distributed systems in association cortex (Larsen and Luna, 2018). Both myelin-sensitive imaging and post-mortem histology have provided evidence for continuous remodeling of intracortical myelin content into adulthood (Grydeland et al., 2019; Hill et al., 2018; Hughes et al., 2018), but it remains unclear whether cortical areas exhibit graded differences from childhood through early adulthood in the developmental timing of myelination according to their position along the S-A axis.

Advances in non-invasive myelin mapping methods using magnetic resonance imaging (MRI) have facilitated investigations of *in vivo* myelin content and cortical microstructure across the human lifespan (Arshad et al., 2017; Edwards et al., 2018). Computing a measure of cortical myelin from the ratio of a T1-weighted to a T2-weighted MRI acquisition (T1w/T2w) has emerged as a highly accessible non-invasive measure that correlates with histological measures of cortical myelin content (Glasser et al., 2014; Glasser and Van Essen, 2011). However, prior studies evaluating age-related differences in T1w/T2w myelin content (Grydeland et al., 2019; Norbom et al., 2020) have not accounted for residual radiofrequency transmit field (B1+) biases, which impact T1w/T2w estimates in an age-dependent manner (Glasser et al., 2021; MacLennan et al., 2022) through their relationship to body size. Because studies that neglected B1+ bias field effects may be prone to spurious results, it is critical to characterize age-related differences in T1w/T2w myelin after mitigating these confounds.

Here, we apply high-resolution cortical T1w/T2w myelin mapping methods from the Human Connectome Project in Development (*N*=628, ages 8-21) to characterize the maturational timing of cortical myelination from childhood through early adulthood. We test whether cortical areas exhibit graded differences in the rate of T1w/T2w myelination according to their position along the S-A axis. We use a novel approach for mitigating B1+ confound effects in T1w/T2w myelin mapping that enables us to detect unbiased age-related differences in T1w/T2w myelin (Glasser et al., 2021). We fit Bayesian spline models across 180 cortical areas in each hemisphere (Glasser et al., 2016a) to characterize different properties of age-related changes in T1w/T2w myelin across the cerebral cortex, including estimates of the variance in T1w/T2w myelin explained by age, rate of change, age of peak growth, and degree of nonlinearity in T1w/T2w myelin development from 8 to 21 years old. Together, these data demonstrate rapid myelination in sensorimotor areas and gradual myelination in distributed cortical association areas during adolescence, which may reflect ongoing plasticity underlying complex information processing and psychological functioning.

## METHODS

### Participants

Neuroimaging datasets included 628 typically developing participants aged 8 to 21 years (53.5% female; see **Figure 1A)** who were part of the Human Connectome Project in Development (HCP-D). The HCP-D is a large cross-sectional and longitudinal study aiming to characterize brain connectivity development in a sample approximating the demographics of youth in the United States with respect to race, ethnicity, and socioeconomic status (Elam et al., 2021; Harms et al., 2018; Somerville et al., 2018). Participants in this sample were recruited across four sites: Harvard University, University of California-Los Angeles, University of Minnesota, and Washington University in St. Louis. Exclusion criteria for recruitment included (i) premature birth (< 37 weeks gestation); (ii) serious neurological condition (e.g., stroke, cerebral palsy); (iii) serious endocrine condition (e.g., precocious puberty, untreated growth hormone deficiency); (iv) long term use of immunosuppressants or steroids; (v) history of serious head injury; (vi) hospitalization >2 days for certain physical or psychiatric conditions or substance use; (vii) treatment >12 months for psychiatric conditions; (viii) claustrophobia; or (ix) pregnancy or other contraindications for MRI. Participants provided written informed consent and assent, and parents of participants under 18 years provided written informed consent for their child’s participation. All procedures were approved by a central Institutional Review Board administered at Washington University in St. Louis (IRB #201603135).

**Figure 1.**
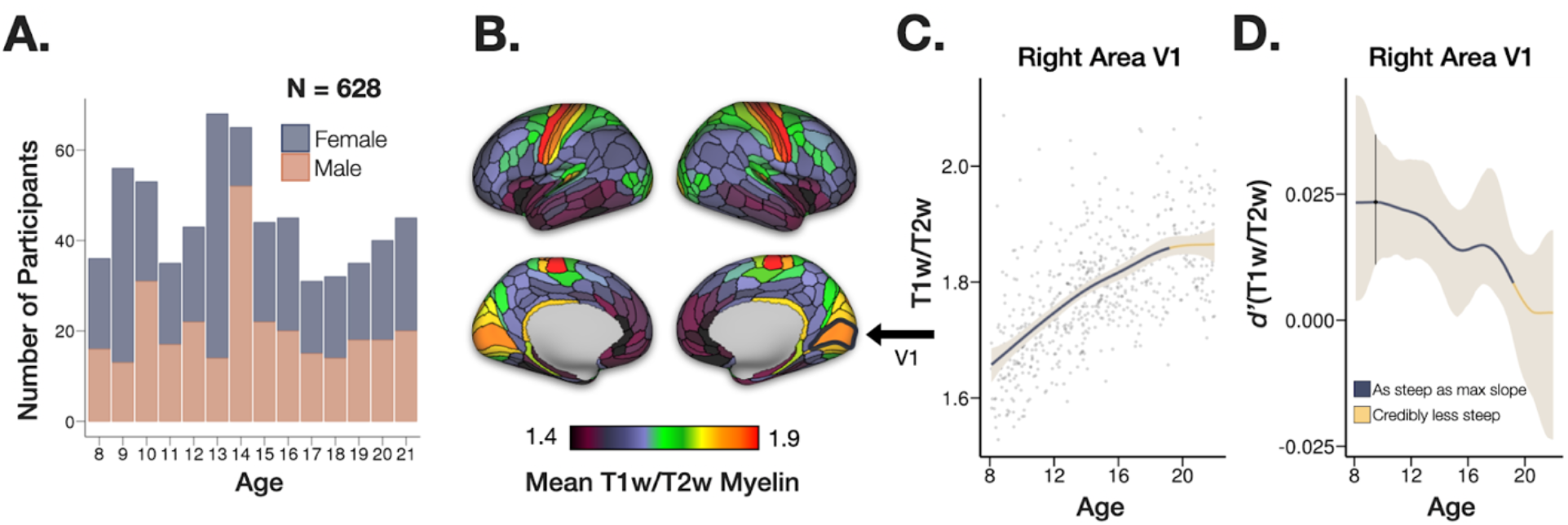
Charting T1w/T2w myelin development during youth. (**A**). Age/sex histogram of the current sample of 628 youth who completed structural neuroimaging as part of the Human Connectome Project-Development (HCP-D). (**B**). T1w/T2w myelin map parcellated using the HCP multimodal atlas (Glasser et al., 2016a) and averaged across participants. T1w/T2w units are arbitrary, representing relative estimates of intracortical myelin content that are comparable within a consistently acquired study. T1w/T2w units of 1.4 and 1.9 correspond to the 2^nd^ and 98^th^ percentiles in this dataset. Data from the right primary visual cortex (highlighted with black border) are shown in panels C and D. (**C**). We fit Bayesian generalized additive models with thin-plate splines to estimate different properties of age-related change in cortical T1w/T2w myelin. Specifically, we estimated the posterior smooth function of T1w/T2w myelin on age for each of the 360 cortical areas (data shown for the right primary visual cortex, highlighted in panel B). The shaded area represents the 95% credible interval of the posterior smooth function. Navy blue segments of the posterior smooth function indicate the slope is credibly as steep as the maximum slope, while gold segments of the posterior smooth function indicate the slope is credibly less steep than the maximum slope (see panel D). (**D**). Bayesian generalized additive models were fitted to estimate the posterior derivative of the smooth function of T1w/T2w myelin on age to characterize the rate of change, with higher values indicating a steeper slope of change per unit age and 0 representing a flat slope (i.e., no age-related change). The shaded area represents the 95% credible interval of posterior derivative estimates. The vertical line marks the age of the maximum median derivative (steepest slope). To identify windows where the rate of age-related T1w/T2w growth credibly slows down, we computed the difference between the posterior derivative of the smooth and the posterior derivative at the age of the maximum median derivative; regions of this 95% credible interval (not shown) that did not include zero were used to mark regions of the smooth that were credibly less steep than the slope at the point of greatest maturation.

This study focuses on cross-sectional data drawn from the publicly available HCP-D “Release 2.0” dataset in the National Institute of Mental Health Data Archive (NDA; N=652). From this dataset, we excluded 21 participants under the age of 8 due to insufficient data available for 5-7 year olds at the time of this study. We excluded 2 additional participants due to face-masking errors during image reconstruction, and we excluded 1 additional participant with poor T1w/T2w data quality indicated by visible artifacts and substantial outliers in T1w/T2w myelin maps, yielding our final sample of 628 youth.

### Image acquisition

High-resolution T1w MRI images were acquired on a 3T Siemens Prisma with a 32 channel head coil using a 3D multi-echo MPRAGE sequence (Mugler and Brookeman, 1990; van der Kouwe et al., 2008) (0.8mm isotropic voxels, TR/TI = 2500/1000 ms, TE = 1.8/3.6/5.4/7.2 ms, flip angle = 8°, in-plane (iPAT) acceleration factor of 2, TA = 8:22, up to 30 reacquired TRs). Structural T2w images were acquired at 0.8mm isotropic using the variable-flip-angle turbo-spin-echo 3D SPACE sequence (Mugler et al., 2000) (TR/TE=3200/564 ms; same in-plane acceleration, TA=6:35, up to 25 reacquired TRs).

Both ‘PreScan Normalized’ (Siemens’ approach for removing the receive coil intensity profile) and non-normalized reconstructions were generated at the scanner. The former were used for image quality review at the scanner, while the latter were as the inputs for subsequent processing (consistent with the use of non-normalized reconstructions as the input for the processing of the HCP Young Adult acquisitions; Van Essen et al., 2013). Both versions (of the T1w) image were used for estimating the B1-receive field to correct for effects of subject motion between the T1w and T2w images. Only the first two echos of the T1w image were used in processing due to artifacts in the later echos that affected surface reconstructions and T1w/T2w myelin maps (Elam et al., 2021).

Volumetric navigators (vNavs) were embedded in the T1w and T2w sequences for prospective motion correction and for selective reacquisition of the lines in k-space that were heavily corrupted by subject motion (Tisdall et al., 2012). Real-time motion correction can substantially reduce bias in brain morphometry analyses, where motion might induce measurable morphometric differences (Reuter et al., 2015; Tisdall et al., 2016). If the T1w or T2w structural scans were nonetheless deemed to be of poor quality at the time of acquisition, they were reacquired (typically immediately in the same imaging session, although sometimes in a different session). Only the single pair of T1w and T2w scans rated the highest in quality were used in subsequent processing. For further neuroimaging protocol description, see Harms et al., 2018. Additionally 2mm isotropic gradient echo and spin echo images were acquired and used for computing the pseudo-transmit field described below (Glasser et al., 2021).

### Image processing

The structural MRI data were analyzed using the HCP Pipelines (Glasser et al., 2013) version 4.0.0, instantiated into the QuNex container environment (qunex.yale.edu), and the data were released as a part of the Lifespan HCP Release 2.0 in the NDA (nda.nih.gov). Briefly, the T1w and T2w volumes were processed through the *PreFreeSurfer* pipeline, which included gradient nonlinearity distortion correction, initial brain-extraction, and rigid registration into an anterior/posterior-commissure aligned ‘native’ space, registration of the T2w volume to the T1w volume using boundary-based registration (BBR; Greve and Fischl, 2009), correction for the receiver coil bias field based on the smoothed square root of the product of the T1w and T2w images, and registration of the structural images to MNI space. Next, the *FreeSurfer* pipeline (v6.0.0; (Dale et al., 1999; Fischl, 2012) optimized for use with high-spatial resolution (greater than 1mm isotropic) structural images was used for computing the ‘white’ and ‘pial’ surfaces, including use of the T2w volume to optimize the pial surface placement. Lastly, the *PostFreeSurfer* pipeline produced cortical surface models in GIFTI format and surface-related data in CIFTI format, and each subject’s cortical surface was then registered to a common 32k_FS_LR mesh using ‘MSMAll’ areal-feature-based cortical surface registration, which is a multimodal registration constrained by cortical myelin maps and resting-state network maps (Glasser et al., 2016b; Robinson et al., 2018, 2014).

Following cortical surface reconstruction, a single researcher with substantial experience vetting structural image data performed a ‘SurfaceQC’ review of the white and gray matter surface placement of each participant’s data, informed by the T1w/T2w myelin maps (Elam et al., 2021; Glasser and Van Essen, 2011). Cases with significant surface placement errors requiring hand editing were flagged for future editing, and excluded from the cohort that has so far been publicly released and thus were not analyzed in this study.

### T1w/T2w myelin map processing

T1w/T2w myelin maps were generated as a cortical ribbon volume and a surface map using the methods described previously (Glasser et al., 2013; Glasser and Van Essen, 2011), together with more recent improvements (Glasser et al., 2021, 2014). Briefly, the T1w/T2w ratio in the voxels between the white and pial surfaces is mapped to the surface mesh in a way that emphasizes the middle layers and underemphasizes voxels near pial and gray/white surface in order to reduce partial volume effects (Glasser et al., 2013). Division of the T1w image by the T2w image mathematically cancels the signal intensity bias related to the sensitivity profile of the radio frequency receiver coils, which is the same in both images in the absence of subject head motion. Taking the ratio will also increase the contrast related to myelin content since both input images show myelin-related contrast (Glasser and Van Essen, 2011), inverted in the T2w image relative to the T1w image. Individual T1w/T2w myelin maps were parcellated according to the HCP-multimodal atlas (Glasser et al., 2016a) using *wb_command-cifti-parcellate* in Connectome Workbench v1.4.2 (Marcus et al., 2011). This computes the average value of all vertices within a parcel and is thus a neuroanatomically constrained form of smoothing. T1w/T2w units are arbitrary, representing relative estimates of intracortical myelin content that are comparable within a consistently acquired study. Further, since the scaling of T1w and T2w images depends on the scanner and acquisition parameters, T1w/T2w units should not be directly compared across studies without appropriate harmonization.

### B1+ transmit field correction of T1w/T2w myelin maps

T1w/T2w myelin maps were initially developed for neuroanatomical analyses such as identifying the anatomical boundaries of cortical areas within individuals (Glasser et al., 2016a) and residual biases from the B1+ radiofrequency transmit field were eliminated using the MyelinMap_BC approach (Glasser et al., 2013); however, this approach also removes genuine low spatial frequency individual differences in the T1w/T2w ratio. Given the increasing interest in using T1w/T2w myelin maps to perform statistical comparisons across individuals and groups with other variables of interest, such as age (Grydeland et al., 2019; Kwon et al., 2018; Norbom et al., 2020), a new approach to B1+ transmit field bias correction was developed (Glasser et al., 2021). B1+ biases are influenced by the loading of the body coil (influenced by both the participant’s head and body). Thus, variables such as head size, body size, and body mass index (BMI), which are often correlated with age and other variables of interest, can modulate B1+ bias and may lead to potentially spurious results in cross-subject statistical analyses of T1w/T2w myelin maps.

In this study, we applied a novel, empirically validated “pseudo-transmit field” correction to mitigate B1+ bias in individual T1w/T2w myelin maps, thereby reducing potentially spurious age-related differences in T1w/T2w myelin content (Glasser et al., 2021). The B1+ correction approach in this study relies on computing a pseudo-transmit field based on the average across phase encoding directions of gradient echo (GRE) images divided by spin echo (SE) images (mimicking the gradient echo T1w scan divided by spin echo T2w scan). First, a reference T1w/T2w myelin map was generated at the group level by finding the scaling between the group average pseudo-transmit field and group average T1w/T2w myelin map that minimizes the correlated left-right differences between the two maps (i.e., the clearly spurious left-right asymmetries). This reference group map was used to correct the individual maps. For the individual correction, the pseudo-transmit map was scaled to minimize the correlated differences between the individual’s T1w/T2w myelin map and the reference T1w/T2w myelin map and the pseudo-transmit map (which includes all differences, not simply left-right ones, and is more robust at the individual level). Prior to estimating this correction, any residual B1-effects due to subject head motion between the T1w and T2w images were also removed using the scanner-computed B1-receive field. The pseudo-transmit field requires regularization by thresholding regions of T2* related signal loss combined with spatial smoothing (with compensation for intensity changes induced by smoothing); it is then scaled to equal 1 at the value where the GRE/SE ratio corresponds to the flip angle prescribed by the scanner, a reference value that is determined at the group level. For more methodological details and validation of the correction, see Glasser et al., (2021), which demonstrates that this correction approach eliminates spurious relationships between the T1w/T2w ratio and B1+ bias modulated by body size.

### Statistical analysis

Bayesian generalized additive models were fitted using the *brms* (Bürkner, 2018) interface to the Stan modeling language (Stan Development Team, 2021). Models were fitted separately to each parcellated cortical area to characterize different properties of age-related changes in T1w/T2w myelin across the cerebral cortex. This approach samples from the posterior smooth function of T1w/T2w myelin conditional on age and covariates, representing age-related changes in the overall concentration of gray matter myelin, and the posterior derivative of the smooth function on age, representing the rate of change across the age range. Importantly, Bayesian thin-plate spline models allow us to estimate both linear and nonlinear age-related changes in T1w/T2w myelin content without specifying a linear or polynomial function *a priori* (Fahrmeir and Kneib, 2011; Hastie and Tibshirani, 2017; Wood, 2017). All statistical analyses were conducted in R version 3.5.1 (R Team, 2018).

We evaluated age-related change in T1w/T2w myelin for each cortical area while controlling for participant sex, scanner, and several covariates related to the B1+ transmit field correction including the scanner transmit voltage, the mean of the pseudotransmit map, four regularization parameters [(1) T2* dropout threshold, (2) smoothing FWHM, (3) correction factor for smoothing’s effect on the pseudotransmit field’s intensities, and (4) the slope parameter of the correction], and a corrected T1w/T2w lateral ventricular CSF regressor (after excluding partial volume voxels and CSF flow effects) that are described in detail in Glasser et al., (2021). The *brms* syntax for the full Bayesian model for each cortical area was expressed as follows:

~~~
brms(T1w/T2w ∼ s(Age, k = −1, bs = ‘tp’) + Sex + Scanner + Transmit_Voltage +
Mean_Pseudo_Transmit_Map + T2_Dropout_Threshold +
Smoothing_FWHM_mm + Smoothing_Correction_Parameter + Correction_Slope + Corrected_CSF_Regressor)
~~~

Thin plate regression splines were used for the smoothing basis, and *k* was left at the default value of −1 which entails setting the basis dimension to 10. We retained the default priors set by *brms*; for regression coefficients this is a flat prior over the reals; for the intercept this is a Student *t* distribution with *df* = 3, *μ* set at the mean of the outcome, and *σ* = 2.5; for the standard deviation of the splines and standard deviation of the error term this is a Student *t* distribution with *df* = 3, *μ* = 0, and *σ* = 2.5. We ran four chains with 4,500 total iterations (2,000 warm-up) for each chain, yielding 10,000 posterior draws in total. The posterior smooth function for each cortical area was used to estimate other properties of age-related change in T1w/T2w myelin, as described below.

### Measurements of age-related change in T1w/T2w myelin

Partial R^2^ values for age splines in each regional model were calculated by taking the difference between full models including an age spline and reduced models without age. These R^2^ values reflect the amount of variance in cortical T1w/T2w myelin content explained by age.

#### Annualized rate of change in T1w/T2w myelin

The annualized rate of change in T1w/T2w myelin was calculated for each cortical area by taking the difference between posterior samples for myelination at age 21 and age 8, and then dividing that difference by the number of years in the age range of the sample. This yielded a posterior distribution for this difference which we use to compute a point estimate using the median and 95% credible interval.

#### Age of peak and slowing T1w/T2w myelin growth

The posterior age of peak growth (i.e., steepest slope) in T1w/T2w myelin was estimated using the *curvish* package in R (Flournoy, 2021). Using the posterior of the smooth function and the method of finite differences at 100 equally spaced points, we evaluate the first derivative. Then, for each posterior sample we find the age at which the derivative is maximized, yielding a posterior density reflecting the probability of the slope being maximized at any given age.

To identify windows in development where the rate of age-related increases in T1w/T2w myelin credibly slowed down, we computed the difference between the posterior derivative of the smooth at each evaluation point and the posterior derivative at the age of maximum median derivative. Regions of this 95% credible interval that did not include zero were used to mark regions of the smooth that were credibly less steep than the slope at the point of greatest maturation.

#### Linearity of age-related changes in T1w/T2w myelin

The smooth function from a generalized additive model was identified as nonlinear when the leave-one-out information criterion (LOOIC; Vehtari et al., 2017) difference between a Bayesian linear and spline model was at least 2.0 times the approximate standard error of its estimate (Bürkner, 2018). This is approximately analogous to a two-sided Z-test. The LOOIC indexes the out-of-sample predictive performance of a model. Therefore, cortical areas for which the generalized additive model fits better are those in which we have credible evidence that this more complex, nonlinear form has greater out-of-sample predictive performance.

The mean absolute posterior second derivative of the smooth function of T1w/T2w on age was calculated as a continuous measure of linearity using the *curvish* package in R (Flournoy, 2021). This measure reflects the rate of change in the slope of age-related T1w/T2w differences.

#### Identifying cortical areas with similar T1w/T2w age-related changes using functional latent clustering analysis

To identify data-driven clusters of cortical areas with similar T1w/T2w smooth functions on age, we applied a functional latent mixture model using the *funHDDC* package in R (Bouveyron and Jacques, 2011; Schmutz et al., 2020). First, for Bayesian models in each cortical area, we subtracted the posterior T1w/T2w myelin estimates at age 8 from predicted myelin values at 500 age bins distributed equally between 8-21 years. This effectively removed intercept differences (i.e., starting myelination level) across cortical areas, which were not of interest in this analysis. Second, we converted a brain region by age matrix into a functional data object for input into the *funHDDC* algorithm. Third, we ran *funHDDC* on the functional T1w/T2w trajectories, allowing the optimal number of clusters to vary freely between 2 and 10. Fourth and finally, we used the *funHDDC::slopeHeuristic* function to identify the most parsimonious model solution based on penalized log-likelihood (Schmutz et al., 2020).

#### Sensitivity analyses

##### 1. Cortical thickness

Prior studies have shown that T1w/T2w myelin and cortical thickness measures are inversely correlated across the cerebral cortex (Glasser et al., 2016a, 2014; Natu et al., 2019) (Glasser et al., 2014; Glasser et al. 2016a; Natu et al., 2019). To demonstrate that our results were specific to age-related differences in T1w/T2w indices of myelin content and are not attributable to concurrent changes in cortical thickness, we fit Bayesian spline models for each cortical area while controlling for areal estimates of cortical thickness (i.e., “`*_V1_MR.thickness.32k_fs_LR.dscalar.nii`”). We then calculated the spatial correlation between parcellated cortical maps of the annualized rate of change in T1w/T2w with and without controlling for cortical thickness.

##### 2. Head motion

Head motion during structural MRI acquisition can impact image quality, inducing spurious artifacts in image intensity and morphometric measurements (Savalia et al., 2017; Tisdall et al., 2016). Age-related differences in motion (younger individuals commonly move more than older individuals) are a common concern when evaluating age-related changes in brain imaging measures (Baum et al., 2018; Reuter et al., 2015; Satterthwaite et al., 2013). Thus, we undertook sensitivity analyses in which participants with the greatest motion estimates were excluded, to test whether the primary results were evident in this lower-motion sample. To do so, we estimated the annualized rate of change in T1w/T2w after excluding 114 participants who exceeded the number of allowed TR reacquisitions for either the T1w (up to 30 allowed) or T2w (up to 25 allowed) acquisition, and thus had a considerable degree of ongoing head motion that was not fully corrected. For these participants, the number of k-space lines that still exceeded the motion threshold after the allowed number of reacquisitions is unknown, and thus the quality of these images is degraded relative to what could have been achieved if there was no limit placed on the number of reacquisitions (Tisdall et al., 2012). We then calculated the spatial correlation between parcellated cortical maps of the annualized rate of change in T1w/T2w with and without including participants with excess head motion.

#### Linking T1w/T2w development with the sensorimotor-association axis of cortical organization

To evaluate whether cortical areas exhibited differences in age-related changes in T1w/T2w myelination according to their position along the S-A axis, we calculated the Spearman correlation between parcellated maps of cortical T1w/T2w myelin development and S-A axis rankings. defined groups of sensorimotor and association areas using the bottom and top quartiles of the archetypal S-A axis delineated by Sydnor and colleagues (Sydnor et al., 2021). We used a spatial permutation test (described in detail below) to assess the significance of spatial correlations between each measure of age-related change in T1w/T2w myelin described above (partial R^2^, rate of change, age of peak growth, degree of nonlinearity) and the S-A axis of cortical organization.

Briefly, S-A rankings were derived from cortical maps of ten fundamental brain features exhibiting systematic variation between lower-order sensorimotor areas and higher-order association areas (Sydnor et al., 2021). These cortical maps measured the anatomical hierarchy from T1w/T2w mapping (Glasser and Van Essen, 2011), the functional hierarchy quantified by the principal gradient of functional connectivity (Margulies et al., 2016), the evolutionary hierarchy quantified by macaque-to-human cortical expansion (Hill et al., 2010), allometric scaling quantified as the relative extent of areal scaling with overall brain size (Reardon et al., 2018), aerobic glycolysis quantified from positron emission tomography measures of oxygen consumption and glucose use (Vaishnavi et al., 2010), cerebral blood flow quantified by arterial spin labeling (Satterthwaite et al., 2014), gene expression quantified by the first principal component of brain-expressed genes (Burt et al., 2018), the first principal component of NeuroSynth meta-analytic decodings (Yarkoni et al., 2011), externopyramidization quantified as the ratio of supragranular pyramidal neuron soma size to infragranular pyramidal neuron soma size (Paquola et al., 2020b), and cortical thickness quantified from structural MRI (Human Connectome Project S1200 data). See (Sydnor et al., 2021) for further details.

#### Spatial permutation test

The statistical significance of spatial correlations (*P*_*spin*_) between parcellated cortical maps of age-related change in T1w/T2w and the archetypal S-A axis were assessed using a conservative parcel-based spatial permutation test that preserves the spatial covariance structure of cortical maps (Alexander-Bloch et al., 2018), as implemented by Váša et al. (2018). This permutation procedure generates a distribution of 10,000 spatial null maps randomly rotated on the spherical cortical surface to retain spatial contiguity and hemispheric symmetry of the original cortical maps. For each parcellated map showing age-related change in T1w/T2w myelin (i.e., R^2^, rate of change, age of peak growth, nonlinearity, latent subgroups), we generated 10,000 randomly rotated “null” maps on the cortical surface. We then estimated the null distribution of Spearman correlation coefficients between the archetypal S-A axis map and the randomly rotated maps of T1w/T2w myelin development. Permutation-based *p*-values (*P*_*spin*_) were calculated based on the number of times the empirical correlation coefficient was higher than the permuted correlation coefficient.

#### False-positive error rate correction

When evaluating statistical significance of spatial correlations between cortical maps of age-related change in T1w/T2w and the archetypal S-A axis, the spatial permutation (“spin”) test provides family-wise control of multiple comparisons (Type I error) by accounting for the spatial and contralateral dependence of areal measurements in cortical maps (Alexander-Bloch et al., 2018). Further, we use the Holm correction (Holm, 1979), which is more powerful than Bonferroni and valid under arbitrary assumptions, to adjust the *p*-values across the 5 tests that are used to test the hypothesis that T1w/T2w development is linked with the S-A axis of cortical organization. The same correction was applied across all 15 spatial correlation tests between pairs of parcellated maps showing different properties of T1w/T2w myelin development and the S-A axis (α=0.05; see Figure 8). For descriptive purposes, we dichotomize regions into those with or without a more complex functional form, and those with or without evident windows where the rate of age-related T1w/T2w growth credibly slows down, comprising 360 tests for each set. We do not attempt to control the family-wise false positive rate for the linearity decisions, or probability of making a sign error (Gelman and Carlin, 2014) when identifying windows where the rate of age-related T1w/T2w growth credibly slows down, as we report these as hypothesis-generating descriptions for which a higher Type I error rate (and lower Type II error rate) is preferable.

## RESULTS

We analyzed high-resolution cortical T1w/T2w myelin maps from 628 participants (ages 8-21 years old; **Figure 1A** and **Figure 1B**) to characterize the development of cortical T1w/T2w myelin content during youth after correcting for transmit field biases (Glasser et al., 2021). The posterior smooth function of T1w/T2w myelin on age and the posterior derivative of the age function were estimated by fitting Bayesian thin-plate spline models across 180 cortical areas in each hemisphere, thereby obtaining measures of the overall effect size, rate, and nonlinearity of age-related change in T1w/T2w myelin across the neocortical sheet. For example, we illustrate age-related changes in T1w/T2w myelin estimated for right area V1 using the posterior smooth function (**Figure 1C**) and the posterior derivative (**Figure 1D**) of T1w/T2w myelin on age.

For a descriptive visualization of cortical myelin maps, we averaged data into three age groups (approximating childhood, mid-adolescence, and emerging adulthood) to illustrate the spatial patterning of age-related increases in T1w/T2w myelin from childhood to adulthood. The pattern of cortical myelination during adolescent development reflects the sensorimotor-association axis reported by Sydnor et al. (2021; **Figure 2** inset right). The sensorimotor-association hierarchy was evident in 8-10 year olds (**Figure 2** left; n=145), with heavily myelinated areas of sensory cortex and lightly myelinated areas of association cortex. This hierarchy was present across all age groups, suggesting that the sensorimotor-association hierarchy in cortical myelin content is reinforced from childhood through adulthood. Relative to younger participants, 14-16 year olds (**Figure 2** middle; n=154) had higher T1w/T2w myelin across sensorimotor and intermediate association areas of the cerebral cortex. In 19-21 years olds, we observed a spatial pattern of cortical myelination extending from primary sensory to multimodal association cortex (**Figure 2** right; n=120).

**Figure 2.**
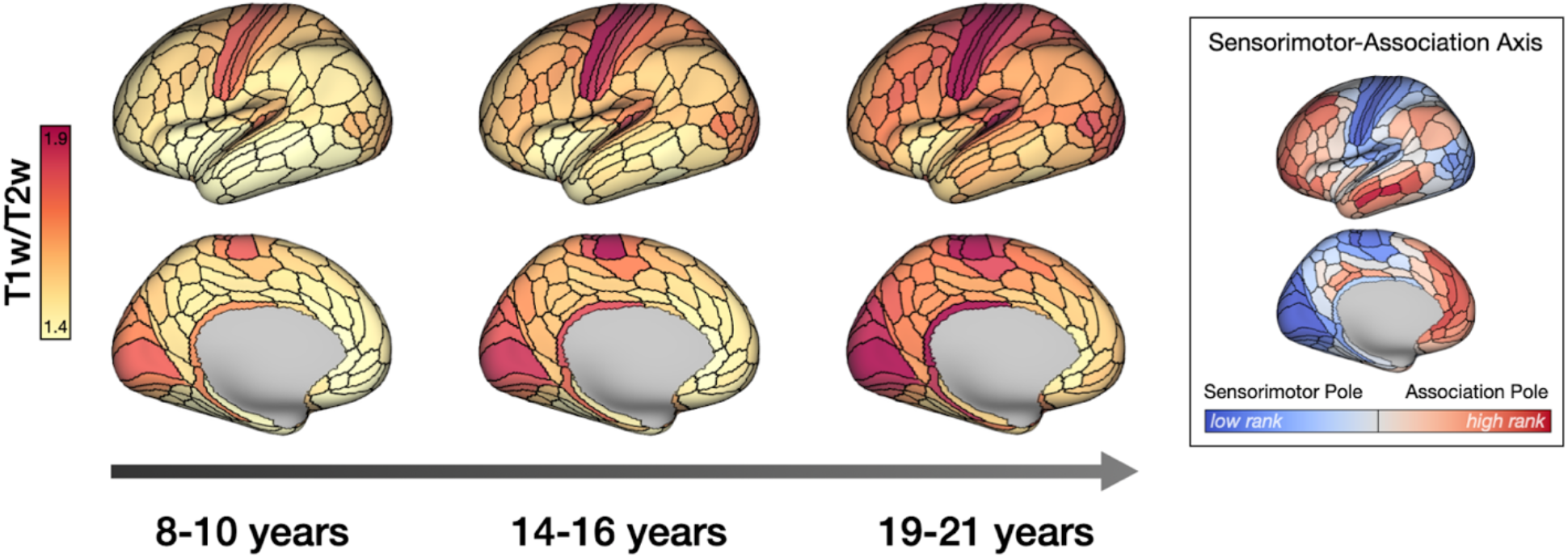
Waves of cortical T1w/T2w myelin development from childhood through early adulthood. Parcellated T1w/T2w myelin maps were averaged across participants within three age groups to illustrate the spatial patterning of age-related increases in T1w/T2w myelin during youth. T1w/T2w units of 1.4 and 1.9 correspond to the 2^nd^ and 98^th^ percentiles in this dataset. Age groups include 8-10 year olds (left; n=145), 14-16 year olds (middle; n=154), and 19-21 year olds (right; n=120). Age-related increases in cortical T1w/T2w myelin were observed across the cortical sheet, reinforcing a sensorimotor-association hierarchy in cortical myelin content. The archetypal sensorimotor-association axis (inset right) was adapted with permission from Sydnor et al. (2021).

Next, we evaluated the topography of the effect size and rate of age-related change in T1w/T2w myelin from childhood through adulthood. We found systematic variation in the amount of age-related variance in T1w/T2w myelin across the cerebral cortex. Specifically, age explained a greater proportion of variance in T1w/T2w myelin in sensorimotor areas such as right area V2 (*R*^2^ = 0.25; **Figure 3B**) compared to association zones where age explained little variance in T1w/T2w myelin, such as the left anterior cingulate cortex (area p32; *R*^2^= 0.028, **Figure 3C**). Figure 3D shows the strong spatial correlation between the variance in T1w/T2w myelin explained by age (*R*^2^) and the cortical area’s position along the archetypal sensorimotor-association axis of cortical organization (*r* = −0.65, *P*_*spin*_ < 0.001).

**Figure 3.**
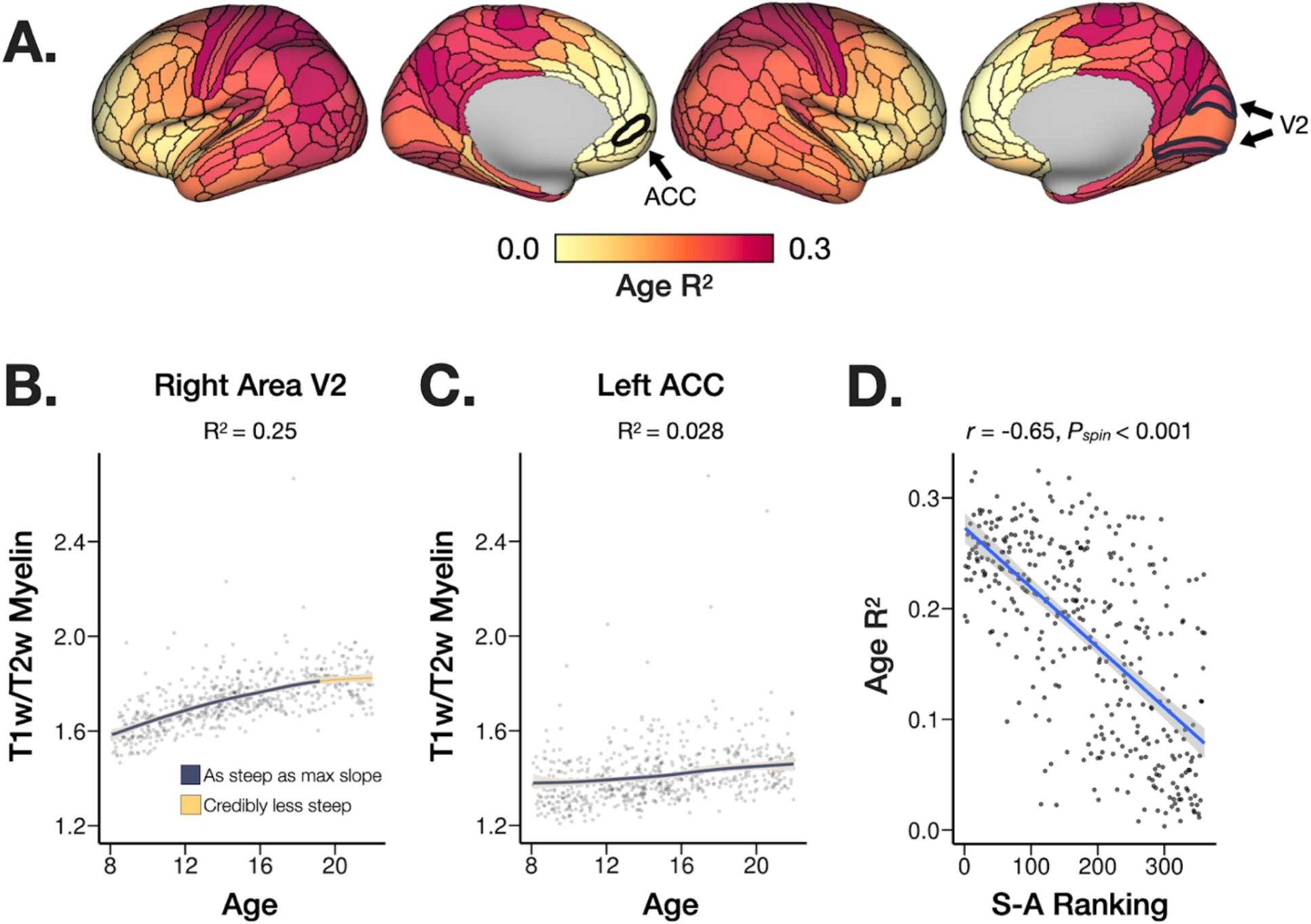
Effect size estimates for regional Bayesian models of T1w/T2w myelin development. (**A**) Partial R^2^ values for age splines were estimated for each cortical area, representing the proportion of variance in T1w/T2w myelin change explained by age. (**B**) Age-related increases in T1w/T2w were relatively strong in sensorimotor areas such as the right area V2 (outlined and labeled in panel A), where age explained 25% of variance in T1w/T2w myelin. (**C**) By contrast, age-related increases in T1w/T2w were relatively weak in heteromodal and paralimbic association areas such as the left anterior cingulate cortex (area p32; outlined and labeled in panel A), where age explained 2.8% of variance in T1w/T2w myelin. (**D**) The variance in T1w/T2w myelin explained by age (R^2^) was correlated with the cortical area’s position along the archetypal sensorimotor-association axis of cortical organization. ACC = anterior cingulate cortex. S-A = sensorimotor-association. *P*_*spin*_ is the permutation-based *p*-value calculated from a conservative parcel-based spatial permutation (“spin”) test.

We also observed spatial variation in the annualized rate of change (AROC) in T1w/T2w myelin estimated for each cortical area (**Figure 4A**). Sensorimotor areas had relatively high AROC in T1w/T2w myelin. **Figure 4B** shows the posterior smooth for the right motor cortex (area 4), an exemplar sensorimotor area with relatively high AROC in T1w/T2w myelin (AROC = 0.025). **Figure 4C** shows the posterior smooth for the left anterior cingulate cortex (area p32), an exemplar association area with relatively low AROC in T1w/T2w myelin (AROC = 0.005). From childhood through early adulthood, spatial variation in the AROC in T1w/T2w myelination reflected S-A hierarchy (**Figure 4D;** *r* = −0.65, *P*_*spin*_ < 0.001), with heavily myelinated sensorimotor areas showing rapid T1w/T2w myelin increase relative to heteromodal and paralimbic association areas, which had slower age-related increases in T1w/T2w myelin.

**Figure 4.**
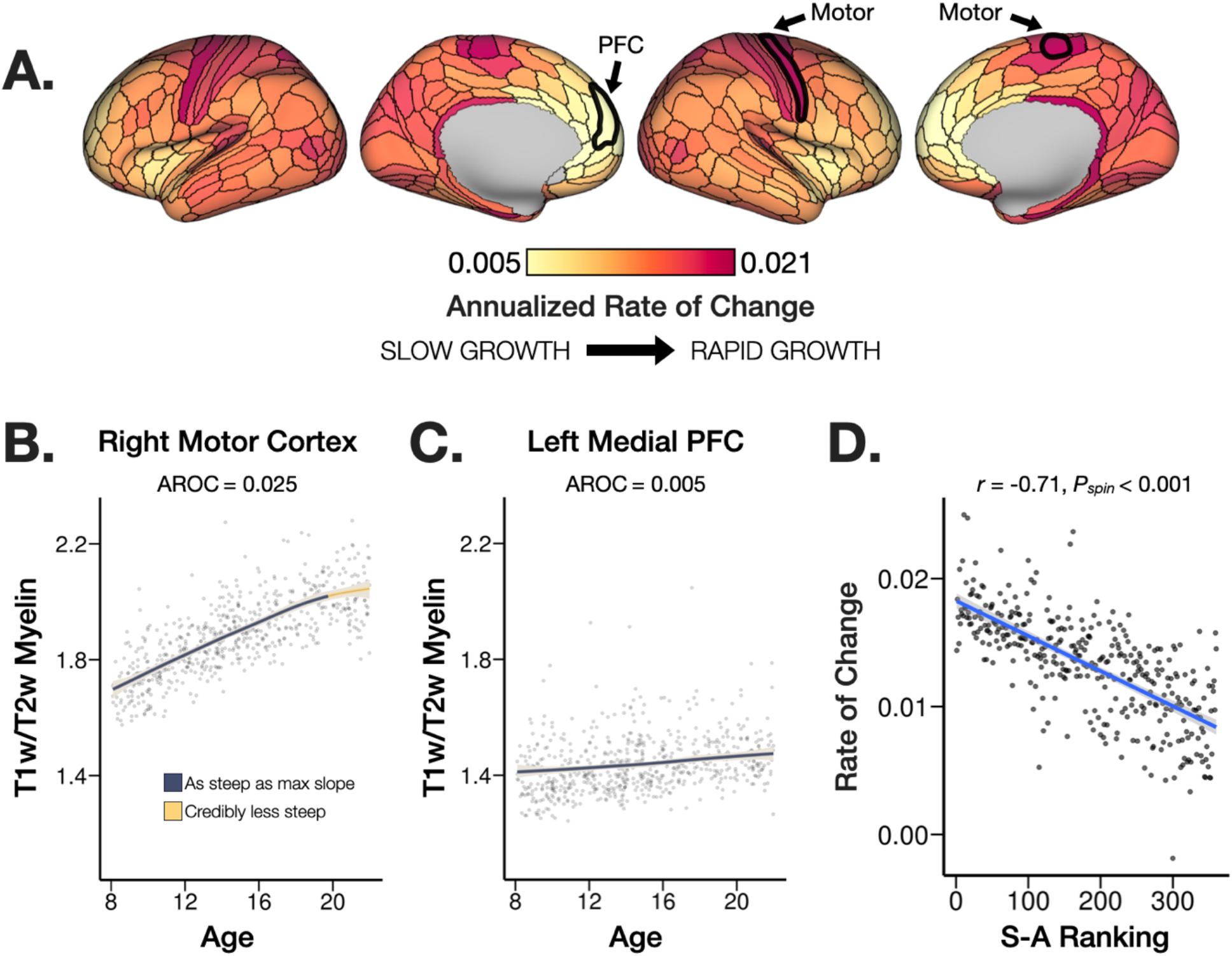
Annualized rate of change in T1w/T2w myelin during youth. (**A**) Annualized rate of change (AROC) in T1w/T2w myelin was estimated for each cortical area. (**B**) The AROC was relatively high in sensorimotor areas such as the right primary motor cortex (area 4; highlighted in panel A, right. (**C**) In contrast, the AROC in T1w/T2w myelin was relatively low in prefrontal and paralimbic association areas such as the left medial prefrontal cortex (area 9m; highlighted in panel A, left). (**D**) The annualized rate of change in T1w/T2w myelin was correlated with the cortical area’s position along the archetypal sensorimotor-association axis of cortical organization. Shaded areas in the plots of panel B and C represent the 95% credible interval of the posterior smooth function on age. Segments of the age curve where the slope is credibly less steep than the maximum slope (indicating a credibly reduced rate of change) are highlighted in gold. PFC = prefrontal cortex. S-A = sensorimotor-association. *P*_*spin*_ is the permutation-based *p*-value calculated from a conservative parcel-based spatial permutation (“spin”) test.

Sensitivity analyses revealed high correspondence between the original parcellated cortical map of annualized rate of change in T1w/T2w myelin and the parcellated map when controlling for cortical thickness in each cortical area (*r* = 0.96, *P*_*spin*_ < 0.001). We also observed a highly consistent spatial pattern of T1w/T2w myelin development when excluding participants with excess head motion (*n* = 451; *r* = 0.97, *P*_*spin*_ < 0.001).

To evaluate whether cortical areas varied systematically in the maturational timing of peak age-related increases in T1w/T2w myelin, **Figure 5A** shows the age of peak T1w/T2w growth based on median posterior age where the first derivative was maximized (i.e. age of steepest slope). **Figure 5B** shows the posterior age of peak T1w/T2w myelin growth for the left area V1, an exemplar sensorimotor area with a relatively early age of peak growth in T1w/T2w myelin (median age = 9.6 years). **Figure 5C** shows the posterior age of peak T1w/T2w myelin growth for the right prefrontal area 8c, an exemplar association area with a relatively late age of peak growth in T1w/T2w myelin (median age = 15.0 years). Spatial variation in the age of peak T1w/T2w myelin growth from 8 to 21 years old closely followed the S-A axis of cortical organization (**Figure 5D;** *r* = 0.60, *P*_*spin*_ < 0.001), with sensorimotor areas showing relatively early peak growth in T1w/T2w myelin compared to heteromodal and paralimbic association areas, which had relatively late peak T1w/T2w myelin growth.

**Figure 5.**
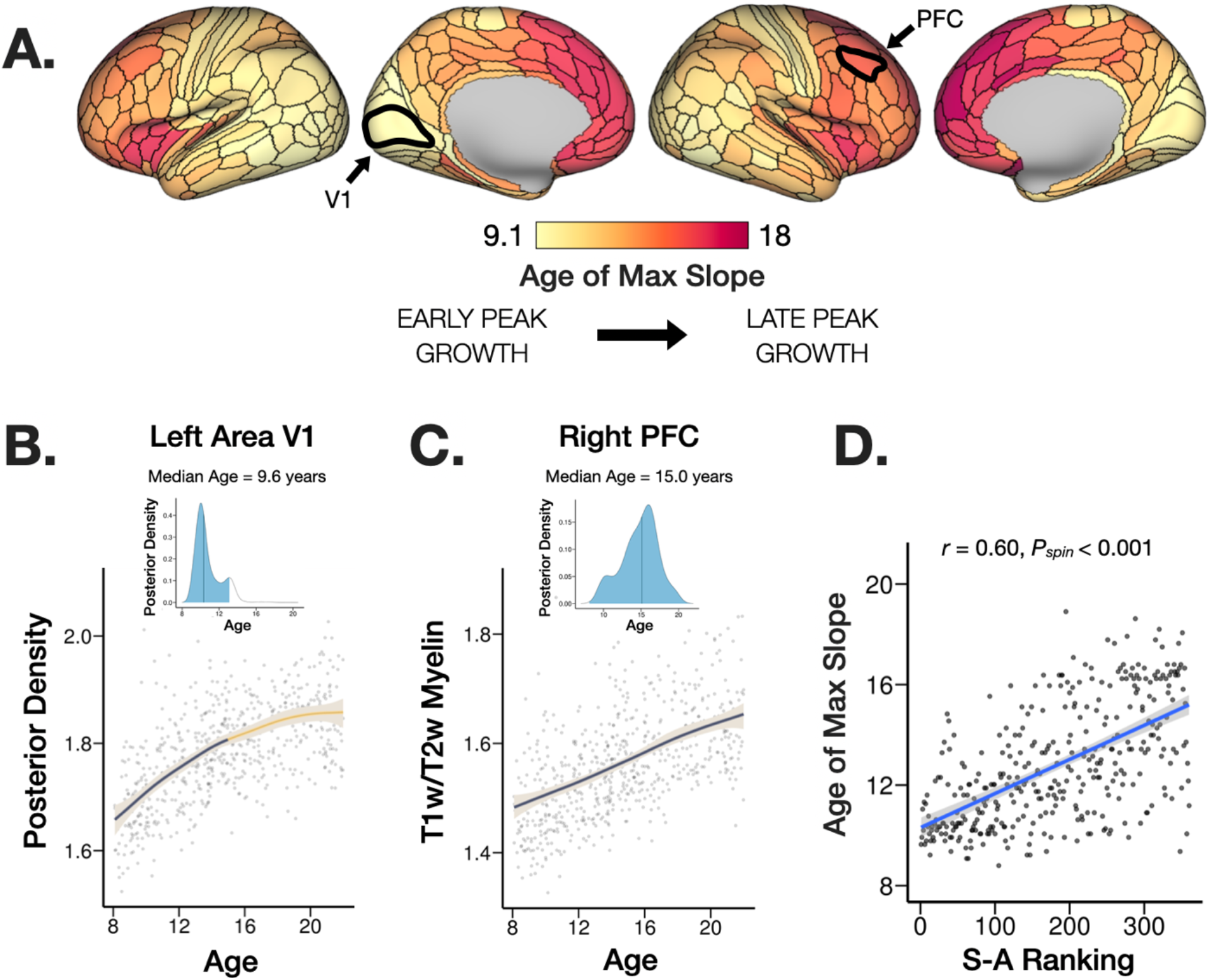
Age of peak T1w/T2w myelin growth during youth. **(A)** The age of peak growth (i.e. steepest slope) in T1w/T2w myelin was estimated for each cortical area as the median posterior age where the slope is at its maximum. (**B**) Sensorimotor areas such as left area V1 (highlighted in panel A) had relatively early age of peak T1w/T2w myelin growth (median age = 9.6 years). (**C**) Association areas such as the right prefrontal cortex (area 8c; highlighted in panel A) had relatively later age of peak T1w/T2w myelin growth (median age = 15.0 years). (**D**) The age of peak T1w/T2w myelin growth was correlated with the cortical area’s position along the archetypal sensorimotor-association axis of cortical organization. Sensorimotor areas exhibited a significantly earlier age of peak growth compared to areas of association cortex. Segments of the age curve where the slope is credibly less steep than the maximum slope (indicating a credibly reduced rate of change) are highlighted in gold. Insets at the top of panels B and C show the posterior density distribution of the age of maximum slope for exemplar areas. The shaded blue area represents the 95% credible interval of the posterior distribution; black vertical lines mark the median of the posterior distribution (values projected on the cortical surface in panel A). PFC = prefrontal cortex. *P*_*spin*_ is the permutation-based *p*-value calculated from a conservative parcel-based spatial permutation (“spin”) test.

Next, we evaluated whether cortical areas varied systematically in the linearity of age-related increases in T1w/T2w myelin, as nonlinear T1w/T2w smooth functions might reflect diminishing age-related changes or a developmental inflection point. **Figure 6A** shows the linearity of age-related changes in T1w/T2w myelin using the mean absolute posterior second derivative, which captures change in the slope over time. **Figure 6B** shows the posterior smooth function of T1w/T2w on age for the right area V1, an exemplar sensorimotor area with nonlinear age-related increases in T1w/T2w myelin (mean absolute second derivative = 0.005). **Figure 6C** shows the posterior smooth function of T1w/T2w on age for the left dorsolateral prefrontal cortex (area p9-46v), an exemplar association area with relatively linear age-related increases in T1w/T2w myelin (mean absolute second derivative = 0.002). Spatial variation in the linearity of age-related increases in T1w/T2w myelin was significantly correlated with the cortical area’s position along the S-A axis (**Figure 6D;** *r* = −0.45, *P*_*spin*_ < 0.001), with sensorimotor areas showing relatively nonlinear T1w/T2w myelin increase relative to heteromodal and paralimbic association areas, which had relatively linear T1w/T2w myelin increase with age. The LOOIC difference between linear and nonlinear spline models indicated that 63 of 360 cortical areas (17.5%) had better out-of-sample predictive performance when including the spline. We found that 116 of 360 (32.2%) cortical areas had segments of the age curve that were credibly less steep than the maximum slope, indicating a credible decrease in the slope of T1w/T2w maturation over time. This deceleration of age-related increases in T1w/T2w myelin occurred primarily between 18 and 21 years old.

**Figure 6.**
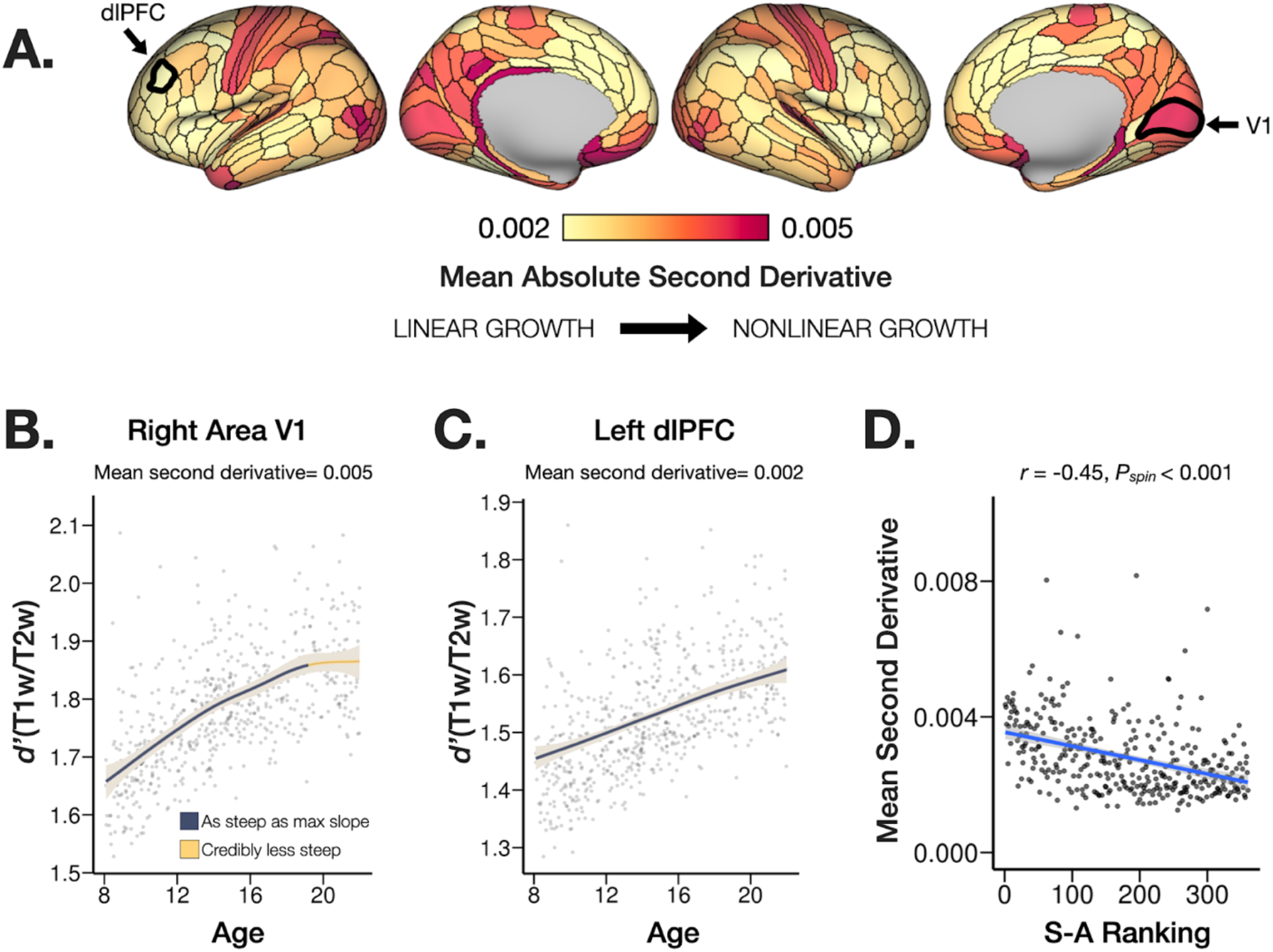
Nonlinearity of age-related changes in T1w/T2w myelin during youth. (**A**) The nonlinearity of age-related changes in T1w/T2w myelin was estimated for each cortical area using the mean absolute posterior second derivative, where higher values indicate more nonlinear growth. (**B**) Sensorimotor areas such as the right area V1 (highlighted in panel A, right) had relatively nonlinear growth, with the posterior derivative decreasing credibly by 19 years old (indicated by gold segment). (**C**) Heteromodal and paralimbic association areas such as the left dorsolateral prefrontal cortex (area p9-46v; highlighted in panel A, left) had relatively linear growth in T1w/T2w myelin (constant slope). (**D**) The nonlinearity of age-related changes in T1w/T2w myelin was correlated with the cortical area’s position along the archetypal sensorimotor-association axis of cortical organization (Sydnor et al., 2021). Sensorimotor areas exhibited significantly higher nonlinearity in age-related increases in T1w/T2w myelin compared to areas of association cortex. Shaded areas in the plots of panel B and C represent the 95% credible interval of the posterior smooth. Segments of the posterior smooth where the slope is credibly less steep than the maximum slope (indicating a credibly reduced rate of change) are highlighted in gold. dlPFC = dorsolateral prefrontal cortex. *P*_*spin*_ is the permutation-based *p*-value calculated from a conservative parcel-based spatial permutation (“spin”) test.

We assessed whether groups of cortical areas demonstrated similar temporal aspects of age-related change in T1w/T2w myelin, as such groups of cortical areas might reflect the development of coordinated patterns of brain connectivity. A functional latent mixture model was used to identify groups of cortical areas exhibiting similar age-related patterns of T1w/T2w myelin growth during youth. The most parsimonious model identified three latent clusters or groups of cortical areas with graded differences in the rate of age-related changes in T1w/T2w myelin (**Figure 7-A**). The spatial patterning of the functional latent clusters closely recapitulated the S-A axis (Sydnor et al., 2021; *F* = 166.6, *p* < 1.0 × 10^−10^): sensorimotor areas at one end of the S-A axis were grouped in latent cluster 1, intermediate association areas were grouped in latent cluster 2, and heteromodal/paralimbic association areas at the opposite end of the S-A axis were grouped in latent cluster 3 based on variation in the developmental timing of T1w/T2w myelination (**Figure 7-B**). Cluster 1 (**Figure 7-C**, magenta) primarily included sensorimotor areas, which had the highest rate of cortical T1w/T2w myelin growth. Cluster 2 (**Figure 7-C**, orange) included heteromodal association areas showing an intermediate rate of cortical T1w/T2w myelin growth. Cluster 3 (**Figure 7-C**, yellow) included paralimbic association areas in medial prefrontal and insular cortex, which had the lowest rate of cortical T1w/T2w myelin growth.

**Figure 7.**
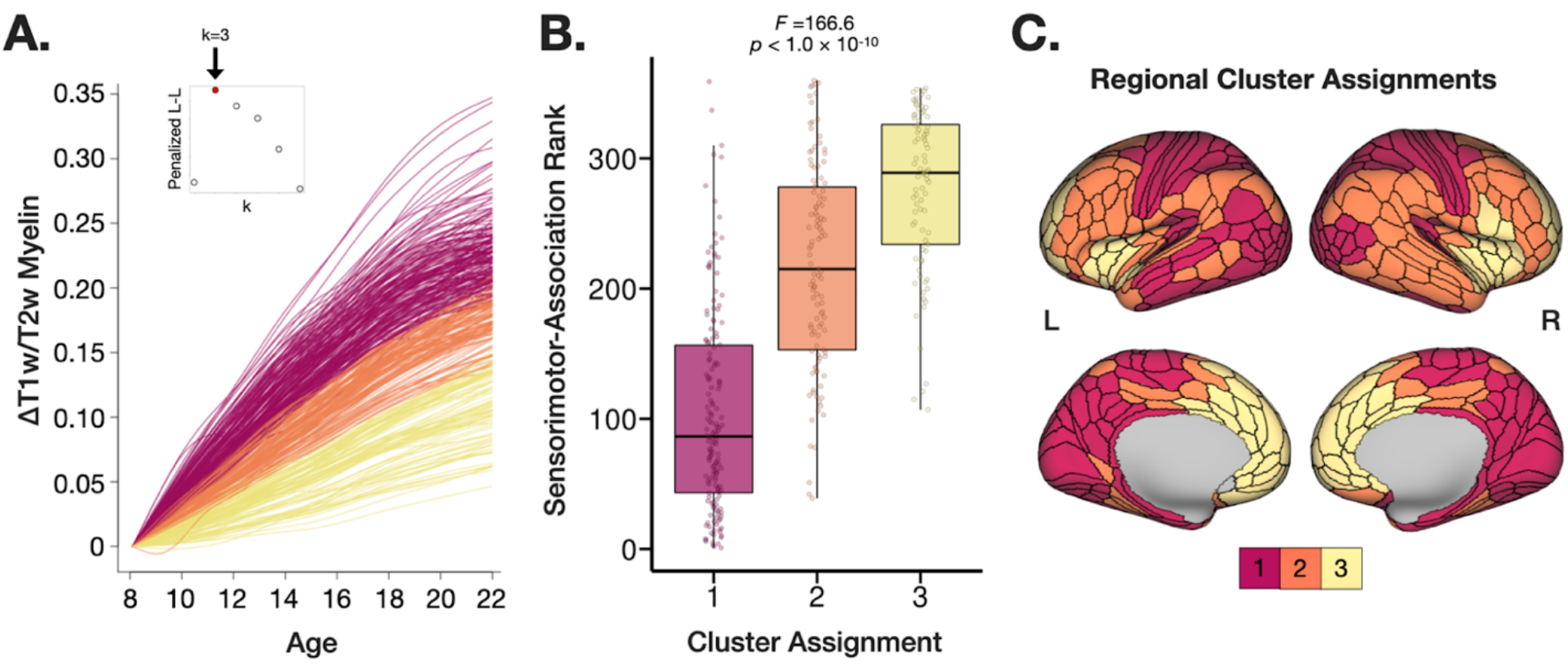
Graded variation across data-driven clusters of cortical T1w/T2w myelin development. A functional latent mixture model was used to identify groups of cortical areas exhibiting similar age-related patterns of T1w/T2w myelin growth during youth. (**A**) The most parsimonious model identified three latent clusters or groups of cortical areas with graded differences in the rate of age-related changes in T1w/T2w myelin. The inset shows the penalized log-likelihood for each model solution from k=2 to k=7 clusters. The best fitting 3-cluster solution is highlighted in red. (**B**) Functional latent clusters closely aligned with the sensorimotor-association axis of cortical organization. Sensorimotor areas, primarily in cluster 1 (magenta), had the highest rate of cortical T1w/T2w myelin growth. We observed graded decreases in the rate of cortical myelin growth in cluster 2 (orange), which included heteromodal association areas in frontoparietal and temporal cortex, and further in cluster 3 (yellow), which included paralimbic association areas in the medial prefrontal and insular cortices. (**C**) Regional cluster assignments based on T1w/T2w myelin development reflect the sensorimotor-association axis. L-L = log-likelihood.

We observed similar spatial patterning across each measurement of age-related changes in T1w/T2w myelin. **Table 1** shows the Spearman correlation coefficients and permutation-based *p*-values (*P*_*spin*_) for each pairwise correlation among measurements. The high correspondence between these measures suggests that different temporal properties of T1w/T2w myelination are yoked during adolescence and depend on the cortical areas position along the S-A axis.

**Table 1.** Correlated properties of T1w/T2w myelin development and cortical Sensorimotor-Association organization. Spatial correlations between parcellated cortical maps highlighting common and dissociable properties of T1w/T2w myelin development and Sensorimotor-Association (S-A) ranking. The lower triangle of the table shows Spearman correlation coefficients and the upper triangle shows uncorrected permutation-based p-values (*P*_*spin*_) calculated from a conservative spatial permutation (“spin”) test. *P*_*spin*_ values from all 15 spatial permutation tests survived family-wise error correction using the Holm method (α = 0.05; Holm, 1979).

|  | S-A Rank | Age R<sup>2</sup> | Rate of Change | Age of Peak Growth | Nonlinearity | Functional Cluster |
| --- | --- | --- | --- | --- | --- | --- |
| S-A Rank |  | < 0.001 | < 0.001 | < 0.001 | < 0.001 | < 0.001 |
| Age R<sup>2</sup> | -0.64 |  | 0.001 | 0.015 | < 0.001 | < 0.001 |
| Rate of Change | -0.74 | 0.86 |  | 0.002 | < 0.001 | < 0.001 |
| Age of Peak Growth | 0.60 | -0.65 | -0.63 |  | < 0.001 | < 0.001 |
| Nonlinearity | -0.45 | 0.47 | 0.57 | -0.47 |  | < 0.001 |
| Functional Cluster | 0.69 | -0.84 | -0.92 | 0.69 | -0.58 |  |

## DISCUSSION

Here we characterize the developmental timing of myelination across the cerebral cortex during adolescence using recent advances in high-resolution T1w/T2w myelin mapping applied to the Human Connectome Project in Development. This study is the first to evaluate the maturational timing of cortical myelination in youth using B1+ transmit field-corrected T1w/T2w myelin maps, providing unbiased estimates of age-related changes in cortical microstructure. We demonstrate graded variation across the cortex in the timing of T1w/T2w myelination during youth, with rapid age-related increases in heavily myelinated sensorimotor cortex, and more gradual age-related increases in lightly myelinated association cortex. This spatial pattern of microstructural brain development recapitulates the sensorimotor-to-association axis of cortical organization and plasticity during ontogeny. Moreover, our findings provide new evidence for protracted cortical myelination in distributed association zones during youth, which could reflect ongoing plasticity underlying complex information processing and psychological functioning.

### Graded variation in cortical T1w/T2w myelination during youth reflects the sensorimotor-association axis of cortical organization

We observed graded variation in the rate and timing of age-related increases in cortical T1w/T2w myelination from 8 to 21 years old. For each cortical area, Bayesian spline models provided posterior estimates of the annualized rate of change, the age of peak T1w/T2w myelin growth, and the degree of nonlinearity in age-related changes. Each of these temporal properties of T1/T2w development followed a cortical topography reflecting sensorimotor-association organization (Sydnor et al., 2021). Sensorimotor areas at one end of the S-A axis had a high rate of T1w/T2w myelination, with early peak growth and relatively nonlinear age-related increases in T1w/T2w myelin. Heteromodal and paralimbic association areas at the opposing end of the S-A axis had a slower rate of T1w/T2w myelination, with late peak growth and relatively linear age-related increases in T1w/T2w. This coupling between measurements of age-related change in T1w/T2w myelin suggest that the myelination of cortical areas follows distinct developmental programs depending on the area’s position along the S-A axis of cortical organization.

A functional latent mixture modeling procedure provided data-driven validation of the systematic variation we observed in T1w/T2w myelin development. The most parsimonious clustering solution identified three clusters of cortical areas based on the rate of age-related changes in T1w/T2w myelin. This approach confirmed that the rate of T1w/T2w myelination during youth parallels the areal position along the cortical S-A axis.

The S-A axis represents a major dimension of cortical organization that captures systematic variation in cortical neuroanatomy, functional connectivity, cerebral metabolism, gene expression, and evolutionary expansion of the cortical mantle (Sydnor et al., 2021). The S-A axis is anchored on one end by sensorimotor areas underlying perception and movement, and at the opposing end by heteromodal and paralimbic association areas underlying cognitive control and socioemotional processing. Further, fundamental neurodevelopmental processes are thought to progress along the S-A axis, such that the early maturation of sensorimotor systems scaffolds the development of association areas that integrate more complex information related to attention, inhibitory control, and socioemotional processes (Larsen and Luna, 2018; Sydnor et al., 2021).

Our results provide new *in vivo* evidence aligned with histological studies demonstrating topographic variation in the rate of cortical gray matter myelination during childhood and adulthood (Flechsig, 1901; Yakovlev and Lecours, 1967), underscoring a neurodevelopmental S-A axis. Graded variation in the rate of cortical myelination could reflect cascading sensitive windows for refining increasingly complex cognitive functions (Takesian and Hensch, 2013). Our findings are also consistent with comparative *post mortem* studies demonstrating prolonged myelination in human association cortex relative to chimpanzees (Miller et al., 2012), which could reflect an uniquely extended window of ongoing plasticity in cortical circuits underlying complex socioemotional and cognitive processing (Larsen and Luna, 2018).

Our main findings also converge with prior neuroimaging studies evaluating age-related changes in cortical myelin content using magnetization transfer (MT) imaging, a quantitative myelin-sensitive contrast, which demonstrate relatively late myelination in association cortices during adolescence into the third decade of life (Paquola et al., 2020a; Whitaker et al., 2016; Ziegler et al., 2019). Our findings are also consistent with a recent study evaluating age-related changes in T1w/T2w myelin from 3-21 years old, which identified global increases in myelination across the neocortex (Norbom et al., 2020). Further, a lifespan study of 8-85 year olds characterized an early age of peak T1w/T2w myelin growth in sensorimotor areas, and a later age of peak myelination in association areas (Grydeland et al., 2019). Despite their strengths, these studies did not control for B1+ transmit field bias, which impacts T1w/T2w myelin estimates in a manner that is systematically related to age, sex, and body size (Glasser et al., 2021; MacLennan et al., 2022). Our study extends prior work by rigorously characterizing the pace and timing of cortical myelin development in youth, using sub-millimeter resolution, B1+ bias-corrected T1w/T2w myelin mapping, and mapping data to a multimodal cortical parcellation in a large, demographically diverse sample.

### Spatial pattern of age-related increases in T1w/T2w myelin remain consistent while controlling for cortical thickness

Cortical thickness and myelin content are systematically related, where thinner areas in sensorimotor cortex have higher myelin content, and thicker areas in paralimbic cortex have lower myelin content (but with exceptions such as the thick, heavily myelinated primary motor cortex; Glasser et al., 2014). Previous studies have observed variable rates of cortical thinning across the cortex during early childhood and adolescence (Shaw et al., 2008; Vandekar et al., 2015). Further, age-related changes in cortical thickness have been difficult to distinguish from changes in cortical myelin content under certain circumstances (Natu et al., 2019). In our study, age-related changes in T1w/T2w myelin were not driven by cortical thinning, as we found highly consistent topography of age-related changes in T1w/T2w myelin after controlling for cortical thickness in each area. This indicates that cortical thinning and myelination during youth are dissociable developmental processes.

### Neurobiological mechanisms: The role of cortical myelin in regulating plasticity

While both myelin-sensitive imaging and post-mortem histology have provided evidence for continuous increases in intracortical myelin content into adulthood (Grydeland et al., 2019; Hill et al., 2018; Hughes et al., 2018), the neurobiological mechanisms promoting cortical myelination throughout the lifespan remain largely unclear. Studies in animal models suggest that cortical myelination provides a mechanism to continuously tune the firing properties of neural circuits to support adaptive behavior within an individual’s environment (Makinodan et al., 2012; Mount and Monje, 2017). Axonal myelination enhances neural signaling speed (McDougall et al., 2018) and plays an important role in regulating the timing of sensitive periods in neurocognitive development by providing structural constraints that limit local plasticity (Takesian and Hensch, 2013). Research using animal models has provided strong evidence for the role of myelin-related proteins such as Nogo-A in reducing plasticity by inhibiting neurite outgrowth and dendritic arborization in cortical circuits (Chen et al., 2000; McGee et al., 2005).

The role of cortical myelination in regulating plasticity during early sensitive periods for sensory cortex development has been demonstrated in animal models (Takesian and Hensch, 2013; Toyoizumi et al., 2013). However, the role of cortical myelination in regulating higher-order cognitive development in humans remains poorly understood. Recent work suggests that a large proportion of neocortical myelin ensheathes axons of parvalbumin-positive (PV+) fast-spiking inhibitory neurons (Micheva et al., 2016). Further, the myelination of PV+ interneurons is critical for maintaining excitation-inhibition (E/I) balance in maturing cortical sensory circuits (Benamer et al., 2020). The maturation of PV+ interneurons and E/I balance in neural circuits is a crucial factor regulating the opening and closure of sensitive periods in neurodevelopment (Takesian et al., 2018; Takesian and Hensch, 2013; Toyoizumi et al., 2013). Moreover, we speculate that cortical myelin development could be linked to organizational gradients of parvalbumin gene expression and the modulation of inhibitory circuitry throughout the cerebral cortex (Burt et al., 2018).

### Future directions

While this study characterized age-related changes in cortical T1w/T2w myelin during youth aged 8-21 years, the cross-sectional design precluded inferences on intra-individual changes in cortical myelin content. Although the HCP-D study contains a longitudinal component (Somerville et al., 2018), it was not feasible to incorporate the longitudinal measurements in this investigation due to ongoing data collection and processing. A more comprehensive lifespan approach could help identify milestones of cortical myelin development and senescence, expanding on previous work (Grydeland et al., 2019). The current study provides an important reference dataset characterizing adolescent cortical myelin development that can serve as a benchmark for replication using the longitudinal subset of the HCP-D sample or other developmental longitudinal datasets such as ABCD (Casey et al., 2018). Such efforts would extend our understanding of cortical myelin development to allow us to evaluate the temporal relationships between cortical myelination and improving cognitive abilities. Future work could also investigate how socioeconomic factors, exposure to early life adversity, or other environmental contexts impact the rate of cortical myelination during youth, as well as the multifaceted relationships between developing cortical microstructure, connectivity, and psychological maturation.

## Acknowledgements

Research reported in this publication was supported by grants R24MH108315, R24MH122820, U01MH109589, and U01MH109589-S1 and by the 14 NIH Institutes and Centers that support the NIH Blueprint for Neuroscience Research, by the McDonnell Center for Systems Neuroscience at Washington University, and by the Office of the Provost at Washington University. Portions of this research were carried out at the Harvard Center for Brain Science using instrumentation supported by the NIH Shared Instrumentation Grant Program (S10OD020039). We would like to thank Melanie Grad-Freilich for assistance with data collection and data access, Erin Reid for assistance with neuroimaging data quality assurance, and Valerie Sydnor for helpful discussion and sharing data related to the sensorimotor-association axis.

## Data Access and Availability

Minimally preprocessed structural neuroimaging data from the HCP-D is available for download through the NIMH Data Archive (https://nda.nih.gov/). Parcellated cortical maps presented in this study will be available for download through the BALSA database upon publication. Code for running statistical analyses presented in this study are available on GitHub (https://github.com/gbaum/hcpd_myelin).

